# LipidCruncher: An open-source platform for processing, visualizing, and analyzing lipidomic data

**DOI:** 10.1101/2025.04.28.650893

**Authors:** Hamed Abdi, Yohannes A. Ambaw, Zon Weng Lai, Ritchie Ly, Chandramohan Chitraju, Shubham Singh, Robert V. Farese, Tobias C. Walther

**Affiliations:** Cell Biology Program, Sloan Kettering Institute, New York, NY, USA; mRNA Center of Excellence, Sanofi, Waltham, MA, USA; Howard Hughes Medical Institute, New York, NY, USA

**Keywords:** lipids, lipidomics, sterols, sphingolipids, phospholipids, bioinformatics, computational biology, mass spectrometry, open-source software, scientific software

## Abstract

**Background:** Advances in mass spectrometry (MS)-based lipidomics have led to a significant surge in data volume, underscoring a need for robust tools to efficiently evaluate and visualize these expansive datasets. While numerous software tools have been developed, current workflows are hindered by manual spreadsheet handling and insufficient data quality assessment prior to analysis. Here, we introduce LipidCruncher, an open-source, web-based platform designed to easily process, visualize, and analyze lipidomic data with high efficiency and rigor.

**Results:** LipidCruncher consolidates key steps of the lipidomics analysis workflow, including data standardization, normalization, and stringent quality controls. The platform also provides advanced visualization and analysis tools that are tailored to interrogate lipidomic data and enable detailed and holistic data exploration. To illustrate LipidCruncher’s utility, we analyzed lipidomic data from adipose tissue of mice lacking the triacylglycerol synthesis enzymes DGAT1 and DGAT2.

**Conclusions:** LipidCruncher fills a specific gap in the lipidomics analysis ecosystem by providing an integrated, quality-focused platform that accepts data from multiple sources and complements existing specialized tools. By bridging the critical divide between data generation and biological interpretation, LipidCruncher facilitates rigorous lipidomics analyses to accelerate the translation of complex lipid profiles into biological insights.

## BACKGROUND

Lipids include diverse classes and thousands of lipid species [1, 2], presenting challenges in identifying changes under different physiological conditions. Despite these complexities, quantification and characterization of lipids are essential for dissecting metabolic and signaling pathways involving lipids.

Mass spectrometry (MS) applied to lipid analyses, either via liquid chromatography coupled to mass spectrometry (LC-MS) or “shotgun” approaches, enable collection of large datasets with quantitation of hundreds to thousands of lipid species [3-5]. Analysis of the resultant lipidomic datasets is challenging and involves several steps. First, mass spectra are processed using specialized software (e.g., *LipidSearch* [6], *LipidXplorer* [7], *LipidFinder* [8] and *MS-DIAL* [9]) that assigns specific lipid species to ion current peaks at particular mass-to charge ratios and quantifies their abundance, while utilizing fragmentation spectra of corresponding ions to enhance identification accuracy and structural characterization. These tools turn mass spectra into a structured dataset that can be the starting point for subsequent bioinformatic analyses.

Other software tools (e.g., *lipidr* [10], *LipidSig* [11], *ADViSELipidomics* [12], *MetaboAnalyst* [13], and *LipidOne* [14]) have been developed to bridge the gap from structured lipidomic datasets to biological insights by providing sophisticated features for processing, visualizing, and analyzing lipidomic data. While each of these specialized tools has advantages for advanced analytics, a gap exists for an accessible platform that accepts native data input from major lipidomics platforms, provides quality control tightly integrated with normalization, and offers comprehensive visualization—all optimized for typical lipidomics workflows. This gap is particularly evident for researchers seeking a solution that does not require bioinformatics expertise.

To address these challenges, we developed *LipidCruncher*, an open-source, user-friendly web-based tool that accommodates diverse data formats and streamlines the transition from semi-processed data to lipid-centric data analysis and visualizations. *LipidCruncher* simplifies and standardizes the workflow, integrating a robust framework for data standardization and normalization with rigorous quality checks to ensure the integrity and reliability of the analysis. Further, its suite of interactive visualizations provides an intuitive interface that bridges the gap between complex lipidomic data and biological interpretations.

## RESULTS

### Description of LipidCruncher workflow

The workflow and features of *LipidCruncher* are organized into three modules (**Figure 1**). The following sections summarize the key capabilities of each module; detailed technical specifications and procedures are provided in **Supplementary Methods**.

**Figure 1.**
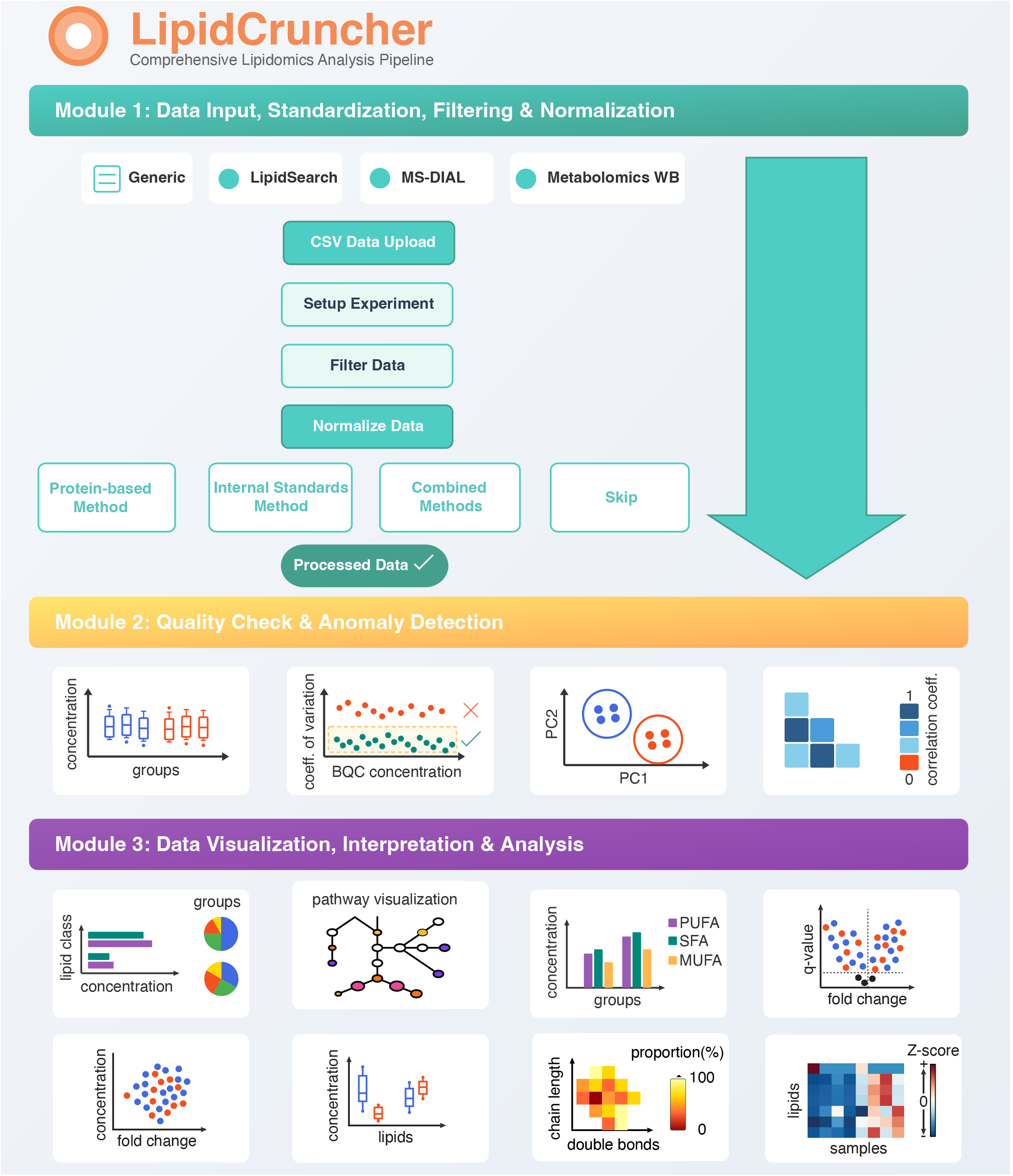
Overview of the *LipidCruncher* analysis pipeline. The workflow is organized into three modules. Module 1 handles data standardization, filtering, and normalization. Module 2 assesses data quality: box plots verify concentration distribution uniformity across replicates, coefficient of variation (CoV) analysis of batch quality control (BQC) samples evaluates measurement precision, and correlation heatmaps and principal component analysis (PCA) enable outlier detection through replicate comparison and sample clustering visualization. Module 3 provides visualization tools: bar and pie charts display lipid class concentration and proportions, metabolic network visualization contextualizes changes across lipid classes, saturation profiles show fatty acid composition (SFA, MUFA, PUFA), volcano plots highlight statistically significant changes with accompanying mean concentration versus fold-change plots and box plots for selected lipids, fatty acid composition heatmaps reveal shifts in chain length and saturation, and comprehensive lipidomic heatmaps with hierarchical clustering display patterns across all detected species.

#### Module 1: Data input, standardization, filtering, and normalization

The first step is data upload (Module 1 of **Figure 1**), for which *LipidCruncher* accommodates outputs from various software platforms. Users can provide data in a generic format (**Supplementary File 1**) or import datasets from *LipidSearch* (**Supplementary File 2**) [6], *MS-DIAL* (**Supplementary File 3**) [9], or *Metabolomics Workbench* (**Supplementary File 4**) [15]. These formats cover widely used platforms in the lipidomics community, including both commercial (*LipidSearch*) and open-source (*MS-DIAL*) mass spectrometry processing software, while also enabling access to publicly available datasets from *Metabolomics Workbench* and providing a generic option for compatibility with any processing pipeline. For the generic format, input data consist of a column for lipid species names accompanied by columns containing abundance values for each sample (**Supplementary File 1**). *Metabolomics Workbench* files follow a similar structure to the generic format but include additional metadata such as sample information and experimental conditions. Input files from *LipidSearch* [6] or *MS-DIAL* [9] contain additional data columns (calculated mass, retention time, quality grades) that enable quality assessment features that are not possible with the generic format.

Upon data upload, *LipidCruncher* standardizes column naming, removes duplicates and invalid entries and replaces null or negative values with zeros. Zero-filtering removes lipid species with excessive missing values across conditions. For *LipidSearch* [6] and *MS-DIAL* [9] inputs, quality-based filtering is also applied. *LipidCruncher* provides four normalization options: (1) no normalization for pre-normalized datasets, (2) internal standard-based normalization, (3) protein-based normalization, or (4) combined normalization using both standards and protein-based methods. For internal standard based normalization, *LipidCruncher* automatically detects common standards including SPLASH LIPIDOMIX® and deuterated labels, or users can upload custom standard lists. Before normalization, the platform generates diagnostics bar plots displaying raw intensity values of each internal standard across all samples. Uniform bar heights indicate consistent sample preparation and instrument performance.

#### Module 2: Quality check and anomaly detection

Assessing the quality of the data is a crucial prerequisite step to processing lipidomic data. After normalization, *LipidCruncher* provides several diagnostic tools to assess data quality (Module 2 of **Figure 1**). Box plots display concentration distributions for each sample, where replicates from the same condition should exhibit similar medians and interquartile ranges. A bar chart quantifies the percentage of zero values per sample to identify anomalies. For datasets with batch quality control (BQC) samples—technical replicates created by pooling equal aliquots from all experimental samples— the coefficient of variation (CoV) is calculated for each lipid species. Users can optionally filter species exceeding a chosen threshold (default 30%). Removed species are displayed, allowing users to selectively restore lipids of biological interest before finalizing.

The dataset is further evaluated using pairwise correlation and principal component (PCA) analyses [16]. Correlation heatmaps reveal deviations from expected patterns that may indicate sample outliers. PCA reduces dataset dimensionality and visualizes replicate clustering, with 95% confidence ellipses drawn around each experimental condition. Samples falling outside these ellipses are flagged as potential outliers. When anomalies are identified, *LipidCruncher* provides the option to exclude aberrant samples.

#### Module 3: Data visualization, interpretation, and analysis

This module provides interactive visualizations for exploring lipidomic data (Module 3 of **Figure 1)**. All plots allow users to hover over data points to reveal more detailed information.

*LipidCruncher* generates bar plots displaying mean lipid class concentrations across conditions with standard deviation error bars and statistical significance indicators. Complementary pie charts illustrate proportional distributions of lipid classes. To facilitate detection of altered metabolic pathways, *LipidCruncher* visualizes lipid abundances in a metabolic network context. The current implementation includes 18 major lipid classes; each one is represented by a circle with diameter corresponding to concentration ratio between conditions and color intensity reflecting the fatty acid saturation ratio (proportion of saturated fatty acids). Separately, fatty acid saturation profiles present the composition of saturated (SFA), mono-unsaturated (MUFA), and poly-unsaturated (PUFA) fatty acids within each lipid class through two complementary views: bar plots of absolute concentrations and stacked bar charts of relative percentages.

Volcano plots display log2 fold-change against statistical significance, highlighting significant changes in the upper and outer quadrants. Users can customize thresholds, filter lipid classes, choose statistical tests, and apply multiple testing corrections. Complementary plots show fold-change versus mean concentration and box plots for selected lipids.

*LipidCruncher* offers two types of heatmaps. Fatty acid composition heatmaps [17] display lipids within a selected class by double bond count (x-axis) and carbon chain length (y-axis). Color intensity indicates each species’ relative abundance within the class. Multiple conditions are shown side-by-side with average double bond and chain length markers overlaid, enabling detection of compositional shifts. Comprehensive lipidomic heatmaps display all lipids (rows) across samples (columns) with z-score color coding. Users can preserve original lipid order or apply Ward’s hierarchical clustering [18] to group lipids with similar concentration patterns.

For statistical comparisons, the platform uses Welch’s t-test for two conditions or Welch’s ANOVA with post-hoc testing for multiple conditions, with options for non-parametric alternatives (Mann-Whitney U, Kruskal-Wallis) [19-22]. P-values are adjusted using Benjamini-Hochberg or Bonferroni methods [23], with a two-level framework controlling false discoveries both across lipid classes and within pairwise comparisons.

For data export, users can download plot data in CSV format or visualizations as SVG files. *LipidCruncher* generates comprehensive PDF reports including all visualizations. While designed for lipidomics, the core processing, quality control, and general visualization features (e.g., bar charts, box plots, PCA, correlation analysis, hierarchical clustering, volcano plots) are applicable to other quantitative omics datasets formatted into the generic CSV input.

### Case study example of utilizing *LipidCruncher* to analyze a lipidomic dataset

#### Data input and data quality analysis for the case study

To demonstrate the workflow and utility of *LipidCruncher*, we analyzed lipidomic data from murine adipose tissue in which the enzymes catalyzing triglyceride synthesis, DGAT1 and DGAT2, were deleted (ADGAT-DKO mice) [24] (**Figure 2A**). The following analysis focuses on illustrating the platform’s capabilities; detailed biological interpretation of this dataset has been reported previously [24]. The study included WT and ADGAT-DKO groups, each with four biological replicates, plus four BQC samples. Lipid analysis was performed using LC-MS/MS, with raw data processed using *LipidSearch* 5.0 [6], and the resultant CSV output files used as input for *LipidCruncher* (**Supplementary File 2**).

**Figure 2.**
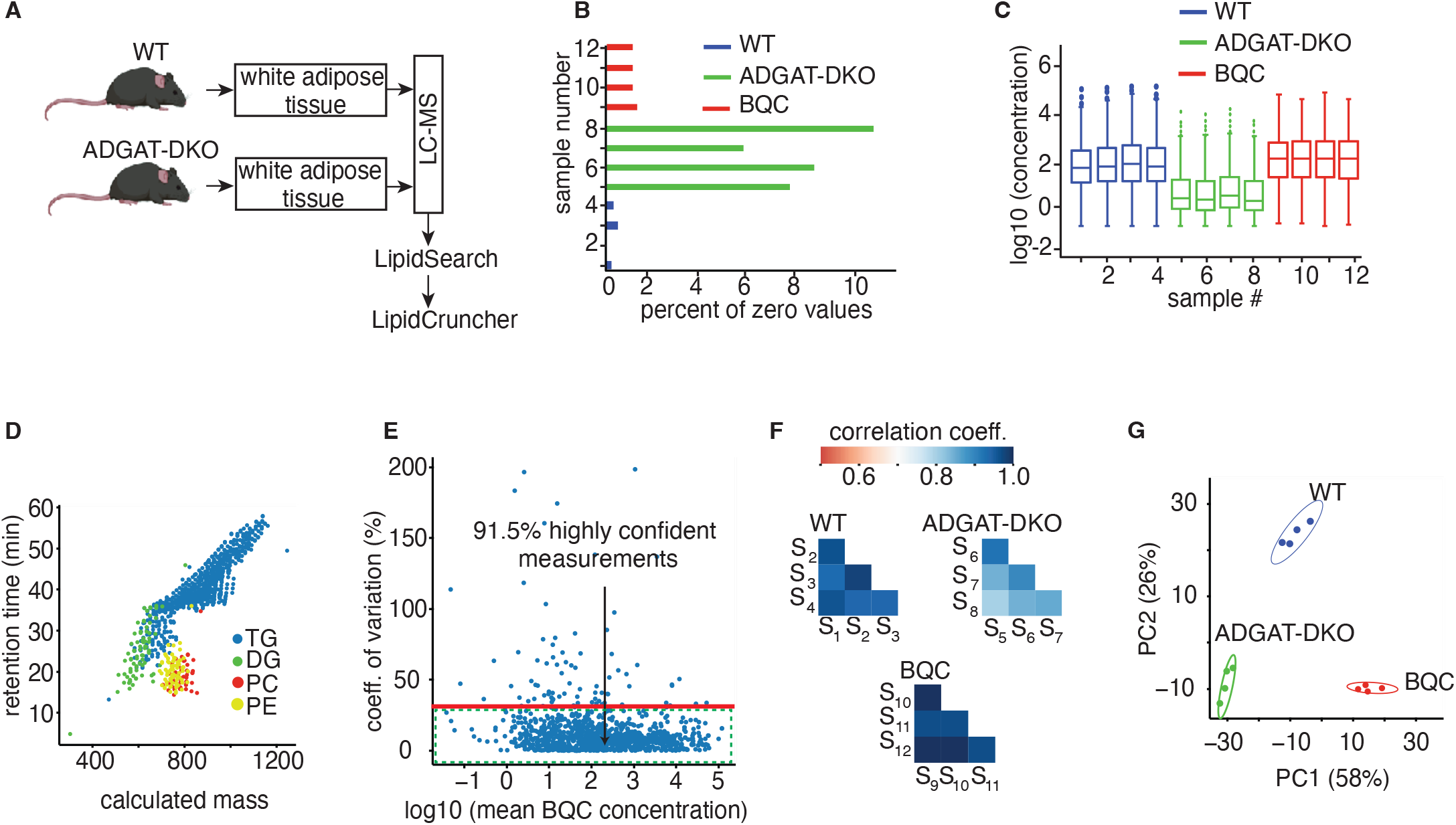
Quality check analysis of lipidomic dataset from ADGAT-DKO mice. A) Experimental design. B) Bar plot showing the percentage of zero values per sample. C) Box plots displaying normalized lipid concentration distributions across samples. D) Scatter plot of retention time versus calculated mass for representative lipid classes (TG, DG, PC, and PE), demonstrating expected clustering patterns based on hydrophobicity. E) Scatter plot of Coefficient of Variation (CoV) against mean concentration for BQC samples. F) Correlation heatmaps showing coefficients between replicates within WT and ADGAT-DKO conditions. G) PCA plot showing clustering of WT and ADGAT-DKO samples with 95% confidence ellipses.

After data standardization and filtering, *LipidCruncher* generated a refined dataset containing 942 endogenous lipid species, spanning 15 lipid classes, along with nine internal standards used for normalization (detailed in **Supplementary Methods**).

Analysis of the dataset revealed a higher proportion of zero values in the ADGAT-DKO samples (∼8%) than WT samples (∼0.4%) (**Figure 2B**), indicating substantial lipidomic changes due to DGAT deletion. Box plots confirmed uniform concentration distributions across replicates within each condition (**Figure 2C)**, validating the normalization process. The retention time versus calculated mass plot demonstrated expected clustering patterns for lipid classes based on their hydrophobicity, with TGs displaying the longest retention times and phospholipids (PC, PE) showing shorter retention times **(Figure 2D)**. This plot serves as a quality check for lipid identification consistency; deviations from expected cluster positions would indicate potential misidentifications.

CoV analysis showed that >90.0% of lipid species had CoV below 30% across BQC samples (**Figure 2E**); species exceeding this threshold were removed. Strong correlations (>0.8) between replicates confirmed data consistency (**Figure 2F**). PCA plots showed distinct clustering within 95% confidence ellipses for each condition (**Figure 2G**), with clear separation between WT and ADGAT-DKO groups reflecting the substantial lipidomic differences. No anomalies were detected during quality assessment.

#### Data visualization, interpretation, and analysis for the case study

Figure 3. represents visualization outputs from *LipidCruncher*. Bar and pie charts (**Figure 3A, 3B)** reveal that TG levels in ADGAT-DKO adipose tissue are reduced by ∼96%, though TG remains the predominant class (decreasing from 98.3% to 77.8% of the total lipid composition). TG depletion is accompanied by increased phospholipid amounts (PC, PE, LPE) and decreased DG and SM levels.

The volcano plot (**Figure 3C**) provides species–level comparison, showing multiple TG species significantly reduced while phospholipid species are elevated. A complementary scatter plot (**Figure 3D**) illustrates the relationship between mean WT concentration and fold-change across lipid classes, revealing reduced TG and DG levels across both high- and low-abundance species. Box plots (**Figure 3E)** enable examination of concentration distributions for individual lipid species across conditions.

**Figure 3.**
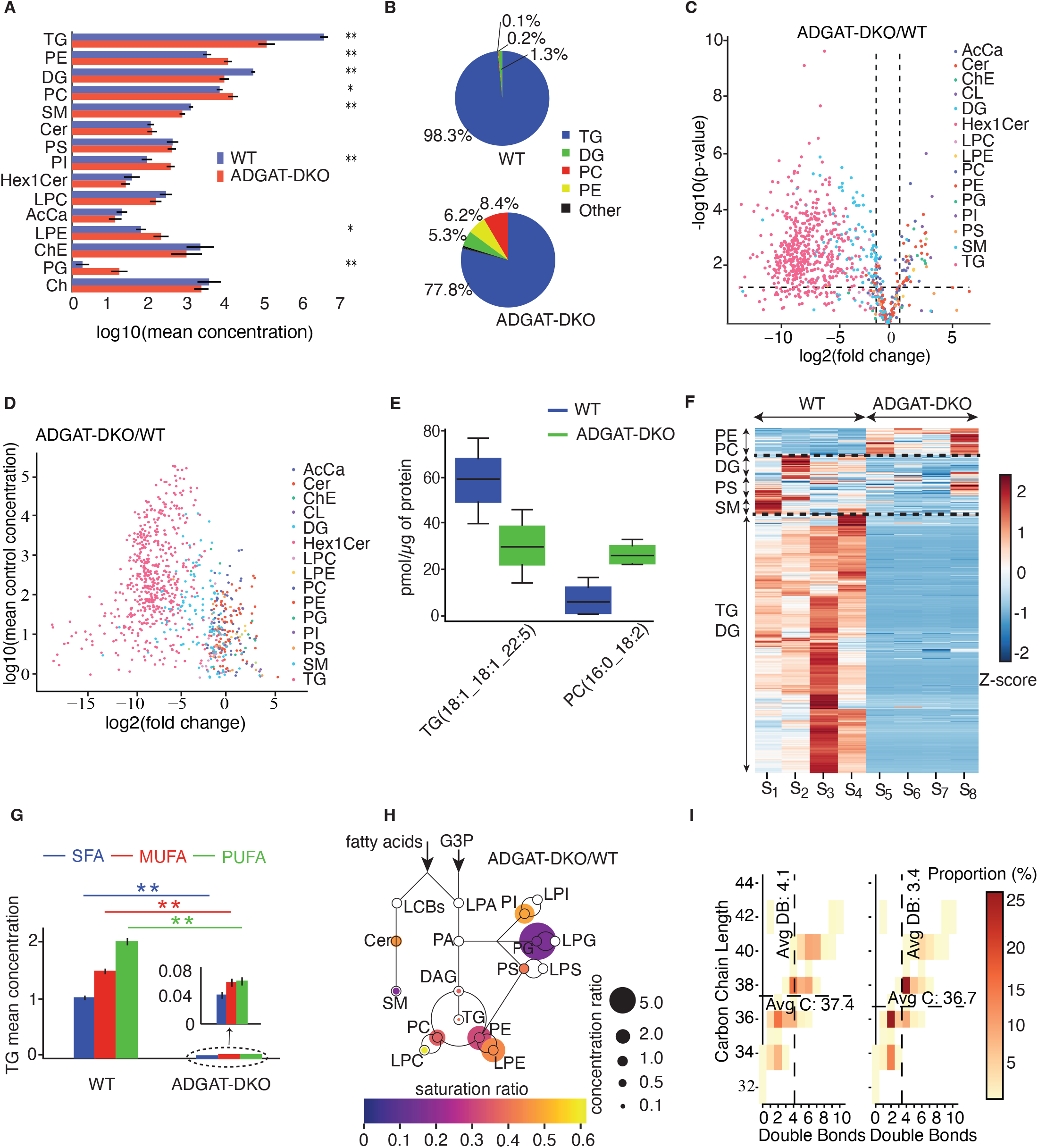
Differential lipid analysis of ADGAT-DKO mice. A) Bar graphs comparing mean lipid class concentrations between WT and ADGAT-DKO mice. Statistical significance: ^*^p<0.05, ^**^p<0.01 and ^***^p<0.001. B) Pie charts showing the proportional distribution of lipid classes. C) Volcano plot displaying log2 fold change versus statistical significance for individual lipid species. D) Scatter plot of mean WT concentration versus fold change. E) Box plot comparing concentration distributions of representative TG and PC species. F) Lipidomic heatmap displaying z-score normalized concentrations for all lipid species, with hierarchical clustering. G) Saturation profile analysis for TG species showing SFA, MUFA and PUFA composition. Statistical significance: ^*^p<0.05, ^**^p<0.01 and ^***^p<0.001. H) Metabolomic pathway visualization showing fold change (circle size) and saturation ratio (circle color). Warmer colors indicate higher saturation; white circles indicate undetected classes. I) Fatty acid composition heatmap for the PE class, displaying lipid distribution by carbon chain length (y-axis) and double bond count (x-axis), with average markers overlaid for each condition.

The lipidomic heatmap (**Figure 3F)** provides a high-resolution view of alterations across 942 lipid species, organized into three clusters: PE and PC lipids, DG, PS and SM species, and TG and DG lipids. Saturation profile analysis of TG species (**Figure 3G)** reveals a shift from PUFAs to MUFAs in the ADGAT-DKO samples, suggesting altered fatty acid metabolism. The pathway visualization (**Figure 3H)** captures dynamic changes in lipid class concentrations, illustrating the interplay between lipid biosynthesis and degradation pathways in the absence of DGAT enzymes.

The fatty acid composition heatmap for PE (**Figure 3I**) reveals a reduction in average double-bond content (4.1 to 3.4) and carbon chain length (37.4 to 36.7) in ADGAT-DKO samples, indicating shifts toward more saturated, shorter fatty chains. These compositional changes reflect altered fatty acid availability driven by impaired TG synthesis.

## IMPLEMENTATION

*LipidCruncher* was developed in Python using Streamlit for the web interface, Plotly and Matplotlib/Seaborn for interactive and static visualizations, and scikit-learn and scipy for statistical analyses and clustering. The platform is deployed on Amazon Web Services (AWS) using Elastic Container Service (ECS), ensuring scalable accessibility. Detailed technical specifications are provided in **Supplementary Methods**.

## CONCLUSIONS

Here we present *LipidCruncher*, an open-source platform that simplifies analysis of lipidomic data by integrating quality assessment, visualization, and statistical tools into a seamless workflow. Designed to accommodate diverse mass spectrometry outputs, it enables researchers to ensure sufficient quality of the data, bypass technical bottlenecks, and focus on extracting biological meaning from complex datasets. The built-in quality control safeguards ensure data integrity, minimizing errors and enhancing reproducibility.

*LipidCruncher* and existing tools serve complementary functions. *MetaboAnalyst* [13] excels in raw spectra processing and advanced multivariate analytics. Among specialized platforms, *lipidr* [10] provides lipid set enrichment analysis, *LipidSig* [11] offers machine learning and network analysis, *ADViSELipidomics* [12] enables batch correction for multi-site studies, and *LipidOne* [14] focuses on building-block level analysis with biomarker discovery. *LipidCruncher* addresses a distinct gap: accessible, integrated analysis of processed lipidomic data optimized for biologists. Rather than attempting to replicate all specialized features, *LipidCruncher* focuses on robust quality control integrated with normalization, comprehensive lipid-specific visualization, and straightforward statistical testing with a two-level multiple testing correction framework. For specialized analyses beyond *LipidCruncher*’s scope—such as pathway enrichment, machine learning, or multi-omics integration—researchers can leverage the dedicated platforms designed for these purposes. This division of capabilities allows each tool to excel at its intended function while collectively serving the broader needs of the lipidomics research community.

Current limitations of *LipidCruncher* merit consideration. While the platform provides multiple quality checkpoints, it cannot identify all potential problems, particularly in upstream processes (peak integration and lipid identification). Regarding lipid identification errors, *LipidCruncher*’s retention time analysis may detect gross misclassifications (e.g., a triacylglycerol incorrectly identified as a phospholipid), but structural misidentifications within the same lipid class (e.g., incorrect fatty acid composition assignments) cannot easily be detected. *LipidCruncher* standardizes lipid nomenclature but does not validate whether reported lipid species are biologically plausible; this responsibility remains with the user. Additionally, while the platform supports outputs from major lipidomics software (*LipidSearch* [6], *MS-DIAL* [9], *Metabolomics Workbench* [15]), it does not directly accommodate all processing tools (e.g., *mzMine* [25] or *Skyline* [26]). Users working with these platforms must standardize their data into the generic CSV format, requiring manual formatting before analysis. Future development will expand format support and incorporate additional quality validation steps.

## Supporting information

guide to supplementary files

## AVAILABILITY AND REQUIREMENTS

**Project name:** LipidCruncher

**Project home page:** https://lipidcruncher.org/

**Source code:** https://github.com/FareseWaltherLab/LipidCruncher

**Operating system(s):** Platform independent (web-based)

**Programming language:** Python 3.9

**Other requirements:** Modern web browser (Chrome, Firefox, Safari, or Edge recommended)

**License:** MIT License

**Any restrictions to use by non-academics:** None

## DECLARATIONS

### Ethics approval and consent to participate

The case study utilized lipidomic data from a previously published study [24]. All animal experiments in that study were performed under the guidelines from Harvard Center for Comparative Medicine. No new animal experiments were conducted for this manuscript. Experimental procedures are detailed in Supplementary Methods.

### Consent for publication

Not applicable.

### Availability of data and materials

The ADGAT-DKO lipidomic datasets analyzed in the case study demonstration are provided as supplementary materials: Supplementary File 1 contains the normalized data in generic format, and Supplementary File 2 contains the raw *LipidSearch* output.

Supplementary Files 3 and 4 contain publicly available third-party datasets included to demonstrate format capability. Supplementary Files 3 is adapted from the MS-DIAL lipidome atlas [28], available under the original publication’s data sharing terms. Supplementary File 4 is from Metabolomics Workbench (Study ID: ST001323) [29], available under Creative Commons Attribution License (CC BY 4.0).

All these datasets are also available on the Farese and Walther lab GitHub site.

### Author contributions

H.A., Y.A., Z.W.L, R.V.F., and T.C.W. conceptualized the application features and design requirements. H.A. developed the application using Python and handled AWS deployment. Y.A. and Z.W.L provided critical guidance on bioinformatics implementation and lipidomic data interpretation across multiple sources. C.C. provided the case study dataset. R.L. and S.S. contributed to scientific discussion and feature suggestions. R.V.F., T.C.W., Y.A., R.L., and H.A. wrote the manuscript. All authors read and edited the manuscript.

## Acknowledgments

We thank members of the Farese & Walther laboratory for suggestions, testing of the software and discussion. We thank Gary Howard for editorial assistance.

## ABBREVIATIONS

BQC: batch quality control
CoV: coefficient of variation
DGAT1: diacylglycerol acyltransferase 1
DGAT2: diacylglycerol acyltransferase 2
LC-MS/MS: liquid chromatography coupled to mass spectrometry
PCA: principal component analysis
WT: wild type
ADGAT-DKO: Adipose-specific DGAT double knockout
SFA: saturated fatty acids
MUFA: mono-unsaturated fatty acids
PUFA: poly-unsaturated fatty acids
PC: phosphatidylcholine
LPC: lysophosphatidylcholine
PE: phosphatidylethanolamine
LPE: lysophosphatidylethanolamine
PI: phosphatidylinositol
LPI: lysophosphatidylinositol
PS: phosphatidylserine
LPS: lysophosphatidylserine
PG: phosphatidylglycerol
LPG: lysophosphatidylglycerol
PA: phosphatidic acid
LPA: lysophosphatidic acid
CL: cardiolipin
CDP-DAG: cytidine diphosphate diacylglycerol
TG: triacylglycerol
DG: diacylglycerol
MAG: monoacylglycerol
Cer: ceramide
SM: sphingomyelin
HexCer: hexosylceramide
CerG1: monoglycosylceramide
CerG2: diglycosylceramide
CerG3: triglycosylceramide
LCB: long-chain base
ChE: cholesteryl ester
CE: cholesteryl ester
Ch: cholesterol
AcCa: acyl carnitine.

Lipid names follow the format Class(chain1_chain2), where each chain is denoted as X:Y, with X representing the number of carbon atoms and Y representing the number of double bonds (e.g., PC(16:0_18:1)). For sphingolipids, the long-chain base is prefixed with ‘d’ (e.g., Cer(d18:1_24:0)). Hydroxyl groups are indicated by semicolon notation (e.g., ;2O). Consolidated format represents total chain composition (e.g., PC(34:1)).

## SUPPLEMENTARY MATERIALS

**Guide to Supplementary Files:** Metadata and guide to supplementary data files. Provides detailed descriptions of file formats, column definitions, sample identifiers, and usage instructions for Supplementary Files 1-4.

**Supplementary File 1:** ADGAT-DKO lipidomic dataset in generic CSV format. Contains normalized lipid concentrations from wild-type and ADGAT-DKO mouse adipose tissue samples used in the case study demonstration.

**Supplementary File 2:** ADGAT-DKO lipidomic dataset in *LipidSearch 5*.*0* format. Contains raw *LipidSearch* output files from the case study experiment, including calculated mass, retention time, quality grades, and intensity values.

**Supplementary File 3:** Example lipidomic dataset in *MS-DIAL* format. Demonstrates *MS-DIAL* compatibility with quality scores, MS/MS matching information, and multi-sample intensity data from an independent experiment.

**Supplementary File 4:** Example lipidomic dataset in *Metabolomics Workbench* format. Demonstrates *Metabolomics Workbench* compatibility with structured sample information and experimental conditions from an independent experiment.

**Supplementary Methods:** Detailed technical procedures for *LipidCruncher* including data format specifications, standardization and cleaning procedures, normalization calculations, quality assessment implementation, statistical analysis framework, visualization implementation details, and experimental procedures for the ADGAT-DKO mouse case study.

