## Supplementary material for "LipidCruncher: An open-source platform for processing, visualizing, and analyzing lipidomic data": guide to supplementary files

*Metadata and guide to supplementary data files for LipidCruncher*

### Overview

This document provides detailed descriptions of the supplementary data files accompanying the *LipidCruncher* manuscript. Files 1 and 2 contain the ADGAT-DKO case study data used in the main manuscript. Files 3 and 4 provide example datasets demonstrating *MS-DIAL* and *Metabolomics Workbench* format compatibility.

| **File** | **Format** | **Description** |
| --- | --- | --- |
| SF1 | Generic CSV | ADGAT-DKO case study (normalized) |
| SF2 | LipidSearch 5.0 | ADGAT-DKO case study (raw) |
| SF3 | MS-DIAL | Mouse adrenal gland fads2 KO vs WT |
| SF4 | Metabolomics Workbench | Mouse HFD serum lipidomics |

### Supplementary File 1: ADGAT-DKO Dataset (Generic Format)

#### Experiment Details

This file contains normalized lipid concentrations from inguinal white adipose tissue (iWAT) of wild-type (WT) and ADGAT-DKO mice [24]. ADGAT-DKO mice lack adipose-specific expression of both diacylglycerol acyltransferase enzymes (DGAT1 and DGAT2), resulting in impaired triacylglycerol synthesis in adipose tissue.

#### Sample Information

| **Sample ID** | **Condition** | **Sample Type** | **Replicates** |
| --- | --- | --- | --- |
| S1–S4 | WT | Biological | 4 |
| S5–S8 | ADGAT-DKO | Biological | 4 |
| S9–S12 | BQC | Technical (pooled) | 4 |

#### Column Definitions

- **Lipids:** Lipid species identifier
- **sample[s1]–sample[s12]:** Normalized lipid concentrations for each sample

*Note: The lipid class (ClassKey) is automatically inferred from the lipid name during LipidCruncher data processing.*

#### Dataset Summary

Total lipid species: 942 endogenous lipids across 15 lipid classes (AcCa, Cer, ChE, CL, DG, HexCer, LPC, LPE, PC, PE, PG, PI, PS, SM, TG).

### Supplementary File 2: ADGAT-DKO Dataset (LipidSearch Format)

This file contains raw *LipidSearch 5.0* output from the ADGAT-DKO case study experiment, including calculated mass, retention time, quality grades, and intensity values. This file demonstrates *LipidCruncher*’s ability to process native *LipidSearch* output with quality assessment features.

#### LipidSearch-Specific Columns

- **CalcMass:** Calculated mass of the lipid species
- **BaseRt:** Base retention time (minutes)
- **TotalGrade:** Quality grade (A = highest, D = lowest confidence)
- **TotalSmpIDRate(%):** Sample identification rate across replicates
- **FAKey:** Fatty acid key with chain composition information
- **MeanArea[s1]–MeanArea[s12]:** Raw intensity values for each sample

### Supplementary File 3: MS-DIAL Format Example

#### Data Source

This dataset is adapted from [28]. The study characterized lipidome diversity across mammalian tissues using *MS-DIAL 4*, a comprehensive software platform for untargeted lipidomics that provides quality scores, MS/MS matching information, retention time, and collision cross-section data. This example file contains lipidomic profiles from mouse adrenal gland tissue comparing fatty acid desaturase 2 (fads2) knockout mice to wild-type controls.

#### Experiment Details

- **Organism:** Mus musculus
- **Tissue:** Adrenal gland
- **Comparison:** fads2 knockout (B6-fads2/J) vs wild-type (C57BL/6J)
- **Diet:** CE2 standard chow

#### Sample Information

| **Condition** | **Genotype** | **Biological Replicates** |
| --- | --- | --- |
| fads2 KO | B6-fads2/J | 3 |
| Wild-type | C57BL/6J | 3 |
| Blank | — | 1 |

#### MS-DIAL Specific Columns

- **Metabolite name:** Lipid species identification
- **Total score:** Composite confidence score (0–100) based on mass accuracy, isotopic pattern, MS/MS match, and retention time
- **MS/MS matched:** TRUE/FALSE indicating MS/MS spectrum validation
- **Average Rt(min):** Average retention time
- **Average Mz:** Average mass-to-charge ratio

#### Data Structure

The file contains both raw intensity values (Height) and normalized concentrations (pmol/mg tissue) in a single export. Raw and normalized data sections are separated by the “Lipid IS” column, which indicates the internal standard class used for normalization. *LipidCruncher* automatically detects this format and prompts users to select which data type to analyze.

### Supplementary File 4: Metabolomics Workbench Format Example

#### Data Source

This dataset is from Metabolomics Workbench [29]. The study examined the effect of high-fat diet on serum lipidome in mice and was conducted at QIMR Berghofer Medical Research Institute.

#### Experimental Design

C57BL/6 mice (n=44) were divided into 4 groups (n=11 per group) and subjected to a 2×2 factorial design with diet (Normal vs. High-fat) and bile acid supplementation (water vs. 1% deoxycholic acid [DCA]) as factors. Treatment duration was 9 months. The study demonstrates the ability of lipidomics to detect both gross changes induced by HFD and specific changes induced by secondary bile acid regulation of liver lipid metabolism.

#### Sample Information

| **Condition** | **Sample IDs** | **n** |
| --- | --- | --- |
| Normal diet + water (Control) | S1A–S11A | 11 |
| Normal diet + DCA | S1B–S11B | 11 |
| High-fat diet + water | S1C–S11C | 11 |
| High-fat diet + DCA | S1D–S11D | 11 |
| Technical QC (TQC) | TQC_1–TQC_12 | 12 |
| Blanks | Blank_1, Blank_2 | 2 |

#### Methodology

- **Organism:** Mus musculus (C57BL/6 strain)
- **Sample type:** Blood serum
- **Lipid extraction:** Matyash method (MTBE-based)
- **Internal standards:** SPLASH LipidoMix + Cer/Sph mixture II (Avanti Polar Lipids)
- **Chromatography:** HILIC (Agilent HILIC Plus RRHD, 100 × 2.1 mm, 1.8 µm)
- **Mass spectrometry:** Agilent 6490 triple quadrupole, ESI mode

#### Lipid Classes Detected

The dataset includes sphingomyelins (SM), ceramides (Cer), and phosphatidylcholines (PC), among other lipid classes.

#### Metabolomics Workbench Reference

Dataset available at: https://www.metabolomicsworkbench.org/ (Study ID: ST001323)

### Usage Instructions

To use these files with *LipidCruncher*:

- Navigate to the *LipidCruncher* web application
- Select the appropriate data format from the dropdown menu
- Upload the corresponding CSV file
- For Metabolomics Workbench format, experimental conditions are automatically detected from the metadata
- For MS-DIAL format, users can select whether to analyze raw or normalized data
- Follow the on-screen prompts to configure the analysis
