## supplementary methods for "LipidCruncher: An open-source platform for processing, visualizing, and analyzing lipidomic data"

### S1. Data Input Format Specifications

#### S1.1 Generic Format

For the generic format, input data are organized with a dedicated column for lipid species classification and structure, accompanied by multiple columns containing values that represent the abundance of each lipid species in a particular study sample. The values are derived from the area under the curve of each lipid species' peak in the mass spectrum, reflecting their relative abundances.

For example, in a study with 12 biological samples—four replicates of three biological conditions—the dataset would consist of rows corresponding to distinct lipid species (e.g., PC(16:0_18:2)). The first column lists unique lipid identifiers followed by twelve columns each containing the intensity/abundance values for this lipid detected in each sample. All inputs must be provided as CSV files.

#### S1.2 LipidSearch Format

*LipidSearch* files include the following columns:

- **LipidMolec:** Lipid species identifier
- **MeanArea columns:** Intensity values for each sample
- **CalcMass:** Calculated mass of the lipid species
- **BaseRt:** Base retention time (minutes)
- **TotalGrade:** Quality grade assigned by *LipidSearch* (A, B, C, or D, where A represents the highest confidence and D the lowest)
- **TotalSmpIDRate(%):** Sample identification rate indicating measurement reliability across samples
- **FAKey:** Fatty acid key containing chain composition information

#### S1.3 MS-DIAL Format

*MS-DIAL* exports alignment results with various columns and quality metrics.

**Mandatory columns:**

- **Metabolite name:** Lipid species identification
- **Intensity columns:** Sample-specific intensity values

**Optional columns (used for quality filtering):**

- **Total Score:** Composite confidence score (0-100) based on accurate mass, isotopic pattern, MS/MS spectrum match, and retention time
- **MS/MS matched:** TRUE/FALSE indicating whether MS/MS spectrum validation passed
- **Average Rt(min):** Average retention time
- **Average Mz:** Average mass-to-charge ratio

Note: The lipid class is inferred from the lipid name; no separate class column is required.

**Raw and normalized data in a single file:** *MS-DIAL* allows exporting both raw intensity values and internally normalized data in a single file. In this format, the file structure is: metadata columns, followed by raw intensity columns for each sample, then a "Lipid IS" column, followed by normalized intensity columns for each sample. The "Lipid IS" column contains the internal standard class used for *MS-DIAL*'s internal normalization and serves as a separator between raw and normalized data sections. Users can choose to analyze either the raw data (for normalization within *LipidCruncher*) or the pre-normalized data. When this combined format is detected, *LipidCruncher* prompts users to select which data type to use.

#### S1.4 Metabolomics Workbench Format

*Metabolomics Workbench* files include:

- Sample information rows with experimental conditions
- Structured data section markers (MS_METABOLITE_DATA_START/END)
- Lipid names in the first column
- Intensity values for each sample in subsequent columns

The experimental conditions from the file are parsed and used to suggest the experimental setup in *LipidCruncher*.

### S2. Data Standardization and Filtering Procedures

#### S2.1 Column Standardization

*Applies to: All formats*

Upon data upload, *LipidCruncher* initiates a standardization protocol. The user is prompted to enter essential experimental parameters, including the number of conditions, descriptive labels for each condition (such as 'WT' for wild type), and the count of replicates per condition. Additionally, the user is asked if the data include BQC samples.

During data standardization, *LipidCruncher* ensures uniform column naming to streamline downstream analyses:

- The column containing lipid species information, including class and structural details, is standardized as "LipidMolec"
- Intensity data for each sample are formatted as "intensity[S₁]," "intensity[S₂]," ..., "intensity[Sₙ]," where N corresponds to the total number of samples in the experiment

#### S2.2 Initial Filtering

*Applies to: All formats*

*LipidCruncher* performs a filtering step to ensure the dataset is prepared for analysis. This involves:

- Removing empty rows
- Removing duplicate entries
- Removing invalid lipid names (e.g., empty strings, special characters, or literals like "NaN")
- Converting non-numeric intensity values to numeric format
- Replacing null or negative values with zeros

**Deduplication** is handled differently depending on the data format:

- **LipidSearch:** When multiple entries exist for the same lipid, the entry with the highest TotalSmpIDRate(%) is selected; if tied, the entry with the best quality grade (A > B > C) is retained
- **MS-DIAL:** When duplicates exist, the entry with the highest Total Score is retained; if no score is available, the first occurrence is kept
- **Generic and Metabolomics Workbench:** Duplicate entries based on LipidMolec are removed, keeping the first occurrence

#### S2.3 Quality-Based Filtering

*Applies to: LipidSearch and MS-DIAL formats only*

For generic and Metabolomics Workbench formats, quality-based filtering is not performed as these formats do not include quality metrics.

**LipidSearch:** *LipidSearch* assigns quality grades to each lipid identification: A (highest confidence), B, C, and D (lowest confidence). By default, *LipidCruncher* accepts grades A and B for most lipid classes. For SM (sphingomyelin) and LPC (lysophosphatidylcholine) classes, grade C is also accepted by default due to the typically lower identification confidence for these classes. Users can customize grade acceptance for each lipid class through the interface. For each unique lipid with multiple entries, the entry with the highest TotalSmpIDRate(%) is selected as this indicates the most reliable measurement across samples.

**MS-DIAL:** *MS-DIAL* provides a Total Score (0-100) reflecting identification confidence. *LipidCruncher* offers three predefined filtering levels, and users can adjust the threshold as needed:

- **Strict:** Total Score ≥ 80 AND MS/MS matched = TRUE (recommended for publication-ready data)
- **Moderate (default):** Total Score ≥ 60 (suitable for exploratory analysis)
- **Permissive:** Total Score ≥ 40 (for discovery-phase research)
- **No filtering:** Total Score ≥ 0 (all identifications retained)

The threshold value is fully adjustable, allowing users to set any custom minimum score based on their specific quality requirements.

#### S2.4 Zero-Filtering

*Applies to: All formats*

A zero-filtering step removes lipid species with excessive zero or below-detection intensity values. Users can configure both the detection threshold and the percentage thresholds for filtering.

**Detection Threshold:** Values at or below this threshold are considered "zero." The default is 30,000 for *LipidSearch* format (reflecting typical instrument noise levels) and 0 for all other formats. Users can adjust this based on their instrument's noise floor.

**Percentage Thresholds:**

- **Non-BQC threshold** (adjustable 50–100%, default 75%): A species fails a non-BQC condition if that condition has ≥ this percentage of replicates at or below the detection threshold.
- **BQC threshold** (adjustable 25–100%, default 50%): A species fails the BQC condition if BQC replicates have ≥ this percentage of values at or below the detection threshold. This slider only appears when the experiment includes BQC samples.

**Filtering Logic:** A species is removed if:

- It fails the BQC condition (when BQC samples are present), **OR**
- It fails **all** non-BQC conditions

Equivalently, a species is kept only if it passes both the BQC condition (when present) and at least one non-BQC condition.

Removed species are listed directly after filtering, allowing review of excluded species and ensuring transparency of the filtering process.

#### S2.5 Handling of Zero Values

While *LipidCruncher* generally does not impute zero values, it interprets zero values as analytes below the detection threshold of the mass spectrometer, representing negligible abundance.

**Exception for log-transformed analyses:** For statistical analyses requiring log transformation (volcano plots, saturation plots, and abundance bar charts), zero values are replaced with one-tenth of the smallest non-zero value in the dataset. This imputation enables the mathematical computation of logarithms while preserving the biological interpretation that these values represent quantities below the instrument's detection limit rather than true absence of the lipid.

### S3. Internal Standard Detection and Normalization

#### S3.1 Automatic Standard Detection

*LipidCruncher* automatically identifies and segregates internal standards based on the following nomenclature patterns:

**SPLASH LIPIDOMIX® Mass Spec Standard** (Avanti Polar Lipids, Cat# 330707-1): Example: DG(15:0_18:1)+D7(s) is recognized as a standard for the DG class.

**Deuterated labels:** Patterns recognized include (d5), (d7), (d9), alternative formats such as +D# notation (e.g., +D7), suffix patterns (_IS, :(s)), and ISTD markers in metadata fields. Standards are identified using pattern matching against lipid names.

#### S3.2 Custom Standards

If users employ alternative standards with different naming conventions, they can upload a CSV file specifying the standards utilized in their samples. Users must first indicate whether the standards exist within their main dataset:

**Mode 1: Extract from Dataset** Select this mode if the custom standards are already present in the uploaded dataset but were not automatically detected. The CSV file should contain:

- A single column with lipid names (must match names in the main dataset)

The system extracts the corresponding intensity values from the main dataset.

**Mode 2: External Standards Data** Select this mode if the standards are external to the main dataset (e.g., separately acquired standard curves). The CSV file should contain:

- Column 1: Lipid names
- Column 2 (optional): ClassKey (lipid class)—if the second column contains non-numeric values, it is treated as ClassKey; otherwise, the class is inferred from the lipid name
- Subsequent columns: Intensity values for each sample (must match the number of samples in the main dataset)

In both modes, column names are automatically standardized, and ClassKey is inferred from lipid names if not explicitly provided.

#### S3.3 Pre-Normalization Verification

Before normalization, *LipidCruncher* organizes the internal standards and generates a separate bar plot for each lipid class, displaying raw intensity values of the internal standards for that class. This enables users to verify consistent sample preparation and instrument performance through uniform bar heights across samples.

#### S3.4 Normalization Calculations

**Internal standard-based normalization:** The platform prompts users to assign the appropriate internal standard to each lipid class and input its concentration. The lipid concentration is calculated by dividing the lipid intensity by the internal standard intensity, then multiplying by the known standard concentration.

**Protein-based normalization:** Users must independently determine protein concentrations (e.g., via BCA assay) and input these values. *LipidCruncher* provides two methods for entering protein concentrations:

- **Manual entry:** Users enter the protein concentration for each sample individually through the interface
- **File upload:** Users upload a CSV file containing a single column named "Concentration" with protein values for each sample in order

The lipid concentration is calculated by dividing the lipid intensity by the protein concentration.

**Combined normalization:** The lipid concentration is calculated by first dividing the lipid intensity by the internal standard intensity, multiplying by the known standard concentration, and then dividing by the protein concentration.

#### S3.5 Post-Normalization Column Naming

After normalization, the column names are updated to reflect the transformed data. The intensity columns ("intensity[S₁]," "intensity[S₂]," ..., "intensity[Sₙ]") are renamed to "concentration[S₁]," "concentration[S₂]," ..., "concentration[Sₙ]" to indicate that the values now represent normalized concentrations rather than raw intensities.

### S4. Quality Assessment Implementation

#### S4.1 Coefficient of Variation Calculation

The coefficient of variation (CoV) is calculated for each lipid species across BQC samples by dividing the sample standard deviation by the mean and multiplying by 100 to express as a percentage. The calculation includes zero values. The mean is log10-transformed for visualization in the CoV scatter plot.

#### S4.2 Coefficient of Variation Filtering

The software provides users with the option to filter data based on CoV values. If users choose to apply CoV filtering, the default threshold is 30%, though this can be adjusted based on specific needs and experimental requirements. A relatively low CoV indicates high fidelity of the data, reflecting reproducible and reliable measurements throughout the experimental workflow.

**Considerations for threshold-based filtering:** Threshold-based filtering may inadvertently exclude biologically meaningful lipids that exhibit naturally high variability. To mitigate this risk, *LipidCruncher* displays all lipids that would be removed in a reviewable table, allowing users to selectively restore specific species they judge to be of biological interest before finalizing the filter.

**Recommendation:** We recommend running the complete analysis pipeline both with and without the species marked for deletion to evaluate their impact on experimental outcomes.

#### S4.3 Correlation Analysis

For correlation analysis, zero values are excluded from the calculation. Pearson correlation coefficients are computed for all pairwise combinations of replicates. A heatmap of correlation coefficients is generated for all replicates. Notable deviations from expected correlation patterns may suggest potential sample outliers, which could arise from biological variability or procedural inconsistencies.

#### S4.4 Principal Component Analysis

**Preprocessing:** Data are standardized by centering each feature to mean of zero and scaling to unit variance. This handles the wide dynamic range of lipid concentrations.

**Dimensionality reduction:** PCA extracts the first two principal components that capture the greatest variance in the dataset.

**Confidence ellipses:** 95% confidence ellipses are drawn around each experimental condition. The ellipse dimensions are calculated from the eigenvalues of the covariance matrix of the PC1 and PC2 coordinates, scaled by a factor of 2 (corresponding to approximately 95% confidence). The ellipse orientation is determined by the eigenvectors.

**Outlier identification:** Samples falling outside the 95% confidence ellipses may indicate potential outliers, suggesting anomalies in sample preparation, data acquisition, or inherent biological differences. The PCA visualization serves as a guide for identifying such samples. Users can select and remove suspected outlier samples from the analysis.

### S5. Statistical Analysis Framework

#### S5.1 Test Selection

*LipidCruncher* offers both automated and manual modes for statistical testing.

**Test type options:**

- **Parametric:** Uses Welch's t-test for two-group comparisons (handles unequal variances) or Welch's ANOVA for multi-group comparisons
- **Non-parametric:** Uses Mann-Whitney U test (two-sided) for two-group comparisons or Kruskal-Wallis test for multi-group comparisons
- **Auto:** Automatically selects parametric tests (Welch's t-test for two-group comparisons and Welch's ANOVA for multi-group comparisons) and applies context-dependent multiple testing corrections based on the number of lipid classes and conditions selected (see S5.2)

#### S5.2 Multiple Testing Correction Framework

The platform implements a two-level multiple testing correction system, allowing users to control false discoveries at different levels of the analysis.

**Level 1 correction (across lipid classes):** Controls false discoveries when testing multiple lipid classes simultaneously.

- **Uncorrected:** No adjustment applied; raw p-values are used
- **FDR (Benjamini-Hochberg):** Controls the expected proportion of false discoveries among rejected hypotheses; recommended for exploratory analyses
- **Bonferroni:** Controls the family-wise error rate by multiplying p-values by the number of tests; more conservative, recommended when false positives must be minimized

**Level 2 correction (pairwise comparisons within multi-group tests):** Controls false discoveries when performing post-hoc pairwise comparisons after a significant omnibus test.

- **Uncorrected:** No adjustment to pairwise p-values
- **Tukey's HSD:** Uses Tukey's HSD for parametric tests (which inherently controls family-wise error rate) or Bonferroni-corrected Mann-Whitney U tests for non-parametric tests
- **Bonferroni:** Uses Bonferroni-corrected pairwise comparisons matching the omnibus test type (Welch's t-test for parametric, Mann-Whitney U for non-parametric)

**Auto mode correction behavior:** When auto mode is selected, the correction methods are determined based on the analysis context:

*Level 1 (across lipid classes):*

- Single class selected → Uncorrected (no between-class correction needed)
- Multiple classes selected → FDR (Benjamini-Hochberg)

*Level 2 (pairwise comparisons):*

- Two conditions → Uncorrected (no post-hoc needed; only one comparison)
- Three or more conditions → Tukey's HSD (controls FWER for any number of comparisons)

#### S5.3 Scale-Appropriate Statistics

For the abundance bar chart analysis, statistics are calculated directly on the appropriate scale: linear scale uses mean and standard deviation of untransformed values, while log10 scale uses statistics computed from log10-transformed values. This ensures mathematically accurate error bar representation on each scale.

#### S5.4 Fold Change Calculation

For volcano plots, fold change is calculated using the original (non-transformed) concentration values by dividing the mean of the experimental group by the mean of the control group. The log2 of this ratio is then computed for visualization. Statistical tests are performed on log10-transformed values to improve normality, while fold change is calculated from original values for biological interpretability.

#### S5.5 Interpretation Guidelines

While *LipidCruncher* applies log transformation to improve normality for parametric tests, it does not explicitly verify the normality assumption required by these tests. Non-parametric tests (Mann-Whitney U, Kruskal-Wallis) do not require this assumption and may be preferred when data distributions are uncertain. We recommend interpreting statistical results as exploratory guides rather than definitive conclusions. The automated significance testing provides a systematic approach to identify potentially interesting lipid changes, but researchers should consider these results in the context of biological relevance, magnitude of change, and consistency across related lipid species. We encourage users to verify notable findings with additional experimental approaches and to consider the biological plausibility of results, especially when working with small sample sizes or highly variable measurements.

### S6. Visualization Technical Details

#### S6.1 Hierarchical Clustering

For lipidomic heatmaps, hierarchical clustering is performed using Ward's method with Euclidean distance. Ward's method minimizes the variance within clusters at each step of the agglomeration process. The pairwise Euclidean distances between all lipid species are first calculated, then clusters are iteratively merged based on the minimum increase in total within-cluster variance. Users select the number of clusters through an interactive interface. The dendrogram leaf ordering is used to arrange lipids in the heatmap visualization.

#### S6.2 Z-Score Normalization

For heatmap visualization, z-scores are calculated row-wise (per lipid) by subtracting the row mean and dividing by the row standard deviation. This enables comparison across samples by standardizing each lipid's concentration profile. The color scale is symmetric around zero, with the range set to plus or minus the maximum absolute z-score value to ensure balanced representation of increases and decreases.

#### S6.3 Fatty Acid Composition Heatmaps

Fatty acid composition heatmaps display lipid distribution within a selected class. The x-axis shows the number of double bonds, the y-axis shows total carbon chain length, and color intensity represents the proportion of each species relative to total class abundance (percentage). Average markers are calculated as weighted means, where each lipid's double bond count or chain length is weighted by its proportional abundance in the class.

**S6.4 Saturation Profile Analysis**

**Fatty acid chain parsing:** For each lipid species, individual fatty acid chains are extracted from the lipid name. For example, PC(16:0_18:1) is parsed into two chains: 16:0 and 18:1. The number of double bonds in each chain determines its classification:

- **SFA (Saturated fatty acids):** 0 double bonds (e.g., 16:0, 18:0, 24:0)
- **MUFA (Monounsaturated fatty acids):** 1 double bond (e.g., 16:1, 18:1, 24:1)
- **PUFA (Polyunsaturated fatty acids):** 2 or more double bonds (e.g., 18:2, 20:4, 22:6)

**Weighted contribution calculation:** Each lipid's concentration is multiplied by the proportion of its fatty acid chains in each saturation category. For example, PC(16:0_18:1) at 100 µM contributes 50 µM to SFA (one of two chains is saturated) and 50 µM to MUFA (one of two chains is monounsaturated).

**Class-level aggregation:** Contributions from all lipid species within a class are summed to generate total SFA, MUFA, and PUFA values for that class. Results are displayed as both absolute concentrations (with standard deviation error bars) and relative percentages.

**Handling of consolidated format lipids:** Lipids reported in consolidated format (e.g., PC(36:2) instead of PC(18:1_18:1)) cannot be accurately classified because the individual chain compositions are unknown. For example, PC(36:2) at 100 µM could represent:

- PC(18:0_18:2): 50 µM SFA + 50 µM PUFA
- PC(18:1_18:1): 100 µM MUFA
- PC(16:0_20:2): 50 µM SFA + 50 µM PUFA

The same total composition yields completely different saturation profiles. *LipidCruncher* automatically detects lipids in consolidated format within selected classes and prompts users to either include them (classification based only on total double bonds, which may be inaccurate) or exclude them (accurate classification for remaining lipids, but reduced total abundance). Single-chain lipid classes (e.g., lysophospholipids, cholesteryl esters, monoacylglycerols) are exempted from this check as they inherently contain only one fatty acid chain.

### S7. Software Implementation

#### S7.1 Software Libraries

*LipidCruncher* was built using the following Python libraries:

**Data manipulation:** pandas for DataFrame structure and data handling; numpy for numerical operations and array manipulation.

**Statistical analysis:** scipy.stats for hypothesis testing (t-tests, Mann-Whitney U, ANOVA); statsmodels for multiple testing corrections (Benjamini-Hochberg, Bonferroni) and Tukey's HSD post-hoc tests.

**PCA and clustering:** scikit-learn for PCA implementation with data preprocessing; scipy.cluster.hierarchy for hierarchical clustering using Ward's method; scipy.spatial.distance for distance calculations.

**Visualization:** Plotly for interactive visualizations with hover, zoom, and filtering capabilities; Matplotlib/Seaborn for static publication-quality graphics.

**Web application:** Streamlit for interactive web application framework.

**Report generation:** reportlab for PDF report generation; svglib for converting SVG graphics in PDF exports; PIL (Pillow) for image processing.

#### S7.2 Deployment

*LipidCruncher* was deployed on Amazon Web Services (AWS) using Elastic Container Service (ECS), ensuring scalable and reliable accessibility for researchers regardless of their computational resources or geographical location.

### S8. Best Practices for Experimental Design

For optimal experimental design and reliable results in lipidomics studies, several best practices are suggested:

**Biological replicates.** There should be at least 4–6 biological replicates for each condition to ensure statistical rigor and account for biological variability.

**Batch quality control samples.** The creation of at least three batch quality control (BQC) samples is crucial. BQC samples are technical replicates generated by pooling equal aliquots from each experimental sample and are essential for evaluating measurement consistency and monitoring instrument performance throughout the analysis.

**Normalization strategy.** We advise using both internal standards (spiked-in lipid molecules of known concentration) and protein-based normalization methods when possible, as this combined approach provides the most robust quantification.

### S9. Case Study: Sample Identifiers and Internal Standards

#### S9.1 Sample Identifier Assignment

For the ADGAT-DKO case study, *LipidCruncher* processed the input dataset and assigned sample identifiers as follows:

| **Sample ID** | **Condition** |
| --- | --- |
| S₁–S₄ | WT |
| S₅–S₈ | ADGAT-DKO |
| S₉–S₁₂ | BQC |

#### S9.2 Internal Standards Used

After data standardization and filtering, *LipidCruncher* automatically identified nine internal standards from the SPLASH LIPIDOMIX® mixture:

1. DG(15:0-18:1)+D7(s)
2. LPC(18:1)+D7(s)
3. LPE(18:1)+D7(s)
4. PC(15:0-18:1)+D7(s)
5. PE(15:0-18:1)+D7(s)
6. PG(15:0-18:1)+D7(s)
7. SM(d36:2)+D9(s)
8. ChE(18:1)+D7(s)
9. TG(15:0-15:0-18:1)+D7(s)

Each internal standard was assigned to normalize lipid species of the corresponding class based on structural similarity.

#### S9.3 Dataset Summary

The refined dataset contained 942 endogenous lipid species spanning 15 lipid classes: AcCa, Cer, ChE, CL, DG, HexCer, LPC, LPE, PC, PE, PG, PI, PS, SM, and TG.

### S10. Experimental Procedures for ADGAT-DKO Mouse Case Study

This section provides detailed experimental methods for the generation of lipidomic data from adipose tissue of mice lacking triacylglycerol synthesis enzymes DGAT1 and DGAT2, which was used to demonstrate *LipidCruncher*'s analytical capabilities in the main manuscript.

#### S10.1 Chemicals

The following reagents were purchased from commercial vendors. Acetonitrile, methanol, water (all HPLC/MS grade), chloroform (HPLC grade), ammonium formate, ammonium acetate, formic acid, and acetic acid were from Sigma-Aldrich. SPLASH® LIPIDOMIX® Mass Spec Standard was from Avanti Polar Lipids, Cat# 330707-1EA.

#### S10.2 Mouse Husbandry

Mice lacking triacylglycerol synthesis enzymes DGAT1 and DGAT2 in adipose tissue (ADGAT-DKO mice) were generated as described in [24]. Mice were housed as per the guidelines from Harvard Center for Comparative Medicine. Mice were maintained in a barrier facility at room temperature (22ºC) on a regular 12-h light and 12-h dark cycle. Mice had ad libitum access to food and water. Mice were fed on standard laboratory chow diet (PicoLab® Rodent Diet 20, 5053; less than 4.5% crude fat). Inguinal white adipose tissue of 12-week-old male mice was used for lipid profiling.

#### S10.3 Sample Extraction and LC-MS/MS Analysis

Lipidomic profiling of inguinal white adipose tissue (iWAT) followed established protocols. Approximately 50 mg of iWAT was homogenized in 1 mL of ice-cold phosphate-buffered saline (PBS) using a bead mill homogenizer. Tissue lysates (50 μg) were then transferred to Pyrex glass tubes with polytetrafluoroethylene-lined caps for lipid extraction.

Lipids were extracted using the Folch method [27]. Briefly, 6 mL of ice-cold chloroform:methanol (2:1, v:v) was added to each sample, followed by 1.5 mL of water. To ensure thorough mixing of polar and non-polar phases, the tubes were vortexed vigorously. SPLASH mix internal standards were spiked into each sample before extraction. Protein concentrations were quantified using a bicinchoninic acid assay (Thermo Scientific, 23225, Waltham, MA) to normalize lipid amounts.

After vortexing, samples were centrifuged at 1100 rpm for 30 min at 4°C to facilitate phase separation. The lower organic phase, containing extracted lipids, was carefully transferred to a new glass tube with a sterile glass pipette to ensure minimal disruption of the interphase layer containing cellular debris and precipitated proteins. Solvents were evaporated under a gentle nitrogen stream until complete dryness. The dried lipid extracts were reconstituted in 250 μL of chloroform:methanol (2:1, v:v) and stored at –80°C until further analysis.

For lipid separation and identification, ultra-high-performance liquid chromatography was coupled with tandem mass spectrometry (MS/MS). Lipid extracts were analyzed using a Thermo Acclaim C30 reverse-phase column (2.1 × 250 mm, 3 μm, Thermo Fisher Scientific) maintained at 55°C. The chromatographic system consisted of a Dionex UltiMate 3000 HPLC system coupled to a Q Exactive Orbitrap mass spectrometer (Thermo Fisher Scientific) equipped with a heated electrospray ionization (HESI) probe.

Each sample (5 μL) was analyzed separately under both positive and negative ionization modes. The mobile phase composition was as follows:

- **Mobile phase A:** 60:40 (v/v) water:acetonitrile with 10 mM ammonium formate and 0.1% formic acid
- **Mobile phase B:** 90:10 (v/v) 2-propanol:acetonitrile with 10 mM ammonium formate and 0.1% formic acid

Mass spectrometric analysis was performed in full-scan/data-dependent MS² mode. Full-scan spectra were acquired at a resolution of 70,000 with an automatic gain control target of 1 × 10⁶ and a maximum injection time of 50 ms, covering an m/z range of 133.4–2000. For data-dependent MS², the top 10 most abundant precursor ions from each full scan were selected for fragmentation using a 1.0 Da isolation window and a stepped normalized collision energy of 15, 25, and 35 units. MS² spectra were recorded at a resolution of 17,500 with an automatic gain control target of 2 × 10⁵ and a maximum injection time of 100 ms.

Lipid identification and data processing were performed using *LipidSearch* software (version 5.0 SP, Thermo Fisher Scientific).
